# Evolution of Regulatory Complexity for Cell-Cycle Control

**DOI:** 10.1101/2021.07.28.454145

**Authors:** Samuel H. A. von der Dunk, Berend Snel, Paulien Hogeweg

## Abstract

How complexity arises is a fundamental evolutionary question. Complex gene regulation is thought to arise by the interplay between adaptive and non-adaptive forces at multiple organizational levels. Using a computational model, we investigate how complexity arises in cell-cycle regulation. Starting from the well-known *Caulobacter crescentus* network, we study how cells adapt their cell-cycle behaviour to a gradient of limited nutrient conditions using 10 replicate *in silico* evolution experiments.

We find adaptive expansion of the gene regulatory network: improvement of cell-cycle behaviour allows cells to overcome the inherent cost of complexity. Replicates traverse different evolutionary trajectories leading to distinct eco-evolutionary strategies. In four replicates, cells evolve a generalist strategy to cope with a variety of nutrient levels; in two replicates, different specialist cells evolve for specific nutrient levels; in the remaining four replicates, an intermediate strategy evolves. The generalist and specialist strategies are contingent on the regulatory mechanisms that arise early in evolution, but they are not directly linked to network expansion and overall fitness.

This study shows that functionality of cells depends on the combination of gene regulatory network topology and genome structure. For example, the positions of dosage-sensitive genes are exploited to signal to the regulatory network when replication is completed, forming a *de novo* evolved cell-cycle checkpoint. Complex gene regulation can arise adaptively both from expansion of the regulatory network and from the genomic organization of the elements in this network, demonstrating that to understand complex gene regulation and its evolution, it is necessary to integrate systems that are often studied separately.

## Introduction

The evolution of cellular complexity is a major question in evolutionary biology. Gene regulation plays a key role in many complex cellular features, from metabolic plasticity in generalist bacteria to cell differentiation in multicellular organisms. With increasing genome size, a larger fraction of the coding genome is devoted to regulatory function compared to information processing and metabolism (van Nimwegen, 2006). Thus, for understanding the evolution of complexity, it is essential to understand how gene regulatory networks evolve.

Changes in gene expression have been associated with adaptation and speciation, suggesting that selection is the main architect of gene regulatory networks (Whitehead and Crawford, 2006; Gilad et al., 2006; Fay and Wittkopp, 2008; Zheng et al., 2011). Yet, models have shown that non-adaptive forces also shape the network (Cordero and Hogeweg, 2006; Lynch, 2007). Large networks are selected to promote evolvability (Cuypers and Hogeweg, 2012, 2014) and mutational robustness (Soyer and Bonhoeffer, 2006). Conversely, small networks are selected to limit off-target interactions in the face of gene expression noise (Jenkins and Stekel, 2008, 2010). Increase in complexity of regulatory networks likely results from a combination of adaptive, mutational and stochastic processes whose interplay remains elusive.

Modelling approaches facilitate mechanistic understanding of gene regulatory network evolution, complementing comparative studies on real data which are limited to extant organisms (Romero et al., 2012). So far, models have focused on simple cases where regulatory networks needed to solve fixed functional challenges, thereby potentially overlooking important evolutionary dynamics, such as neutral variation during long periods of stasis (Quayle and Bullock, 2006). To overcome these issues in understanding the evolution of complexity, we turn to modelling of the cell-cycle, which provides an intrinsic and dynamic fitness criterion for a cell’s regulatory network. Even in prokaryotes, cell-cycle control consists in integration of internal and external cues for correct timing of major cellular events such as genome replication, growth and cell division. Furthermore, due to the lack of strictly separated stages as in eukaryotes, replication impacts ongoing gene expression (Pelve et al., 2011; Paijmans et al., 2016; Walker et al., 2016; Jaruszewicz-Błońska and Lipniacki, 2017).

Cell-cycle regulation in prokaryotes is best understood in the model species *Caulobacter crescentus*, an alpha-proteobacterium that lives in nutrient-poor freshwater streams (Sánchez-Osorio et al., 2017). The five main transcription factors that orchestrate the *C. crescentus* cell-cycle have been identified: CtrA, GcrA, DnaA, CcrM and SciP (Tan et al., 2010; Quiñones-Valles et al., 2014). These five genes produce a cyclic expression pattern that drives all downstream cell-cycle functions (Zhou et al., 2015) and which can be reproduced in a simple Boolean model (Quiñones-Valles et al., 2014; Sánchez-Osorio et al., 2017). We use this model as starting point to study the evolution of complex cell-cycle behaviour in cells exposed to more challenging, nutrient-limited conditions.

## Results

### Model

We define a cell-cycle by the five established core genes from *C. crescentus* (dubbed g1–5) whose combined expression states represent the four cell-cycle stages observed in Quiñones-Valles et al. (2014) (dubbed G1, S, G2 and M; see Methods for a more detailed description of the model). To allow evolution of this well-studied system, we embed the Boolean cell-cycle network in cells that die, replicate their genome, and divide. The gene regulatory network is coded in genomes consisting of discrete *beads*, i.e. genes and binding sites (this representation is known as a beads-on-a-string genome; e.g. Crombach and Hogeweg, 2007). Noise is introduced through sequence-specific binding affinities between gene products and binding sites (i.e. inverse of Hamming distance between bitstrings of length 20); out of all expressed genes, only one may bind a given binding site each timestep.

A cell divides once it completes the four-state gene expression cycle upon reaching the division stage (M). A crucial part of the cell-cycle is genome replication, which happens in discrete chunks of *n* beads when a cell is in the replication stage (S). How long a cell takes to replicate its entire genome depends on the genome size *L* and on the nutrient abundance *n* in its environment (i.e. chunk size). The need to execute multiple replication steps in poor nutrient conditions provides a drive for the evolution of more regulation. Yet replication takes time, so there is inherent selection against genome expansion as larger genomes take longer to replicate.

### Initialization

Cells with the *C. crescentus* cell-cycle network (Quiñones-Valles et al., 2014) encoded as desribed above were pre-evolved for 10^5^ timesteps in rich nutrient conditions to adapt to the model formalism and specifically the newly introduced stochasticity in regulatory dynamics. The resulting most common genotype is streamlined and regulates a cell-cycle that is robust to the intrinsic stochasticity (cf. Jenkins and Stekel, 2010, see also “Evolution of *C. crescentus* cell-cycle under stochastic gene expression” in Appendix 1). With this genotype, we set up 10 replicate evolutionary experiments (R1–R10) under nutrient limitations (Fig. 1, S5). Nutrients are increasingly limited along a spatial gradient and also decrease based on local cell density (see Methods). To expand their range to poorer conditions, cells need to evolve to adjust the duration of the replication stage (S) such that the genome can be fully replicated before reaching the division stage (M).

**Figure 1:**
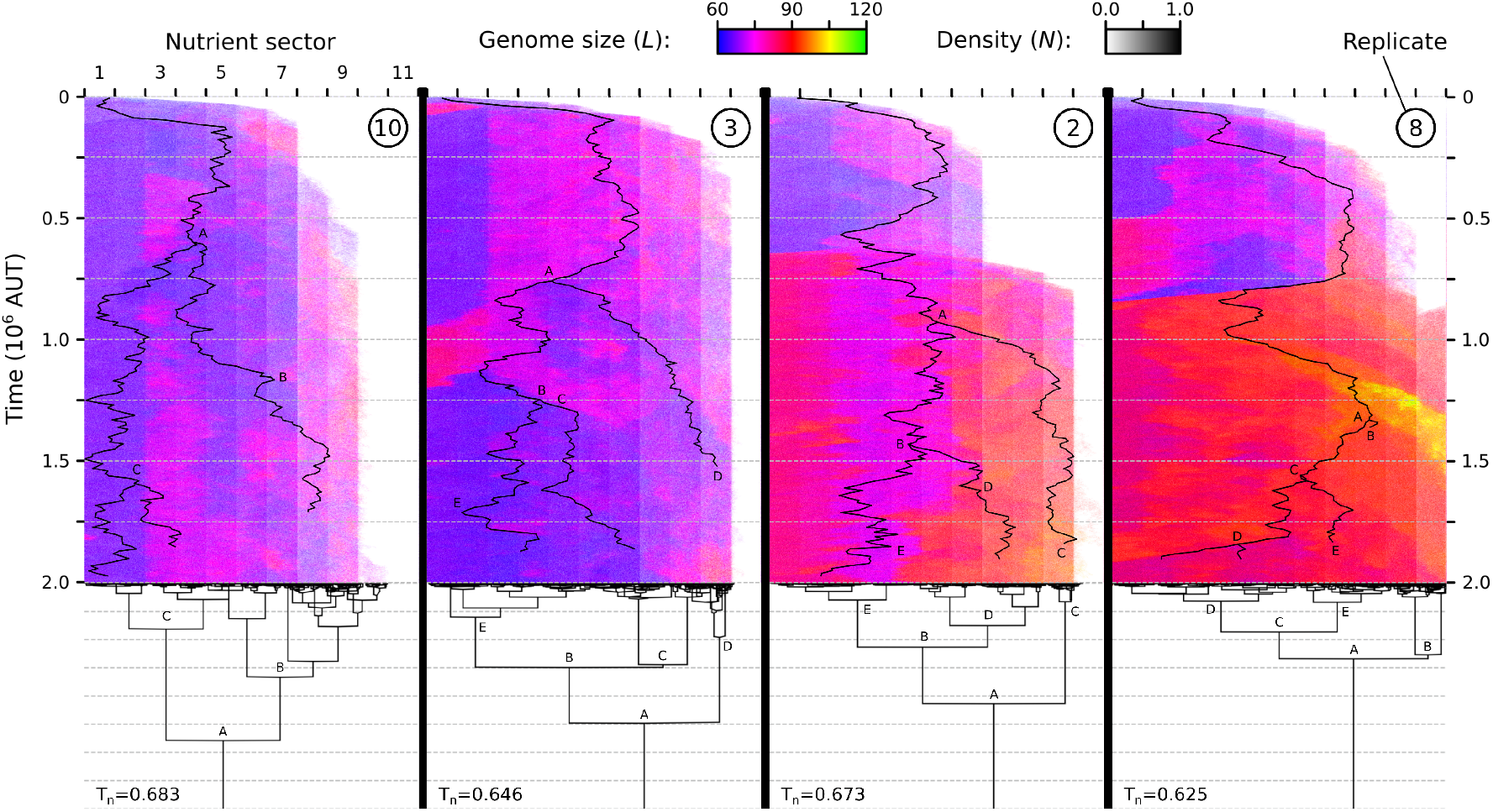
Replicate populations (R1–R10) invade and adapt to a nutrient gradient. Replicates are ordered from smallest to largest final population. Top: space-timeplots with population density and average genome size, bottom: cladogram of the final population at *t* = 2 . 10^6^, including the treeness statistic (*T*_*n*_; see “Evolutionary signatures of generalist and specialist strategies”). For the main ancestral lineages, their location on the gradient is shown on the space-timeplots with letters identifying the corresponding branching events in the cladogram.

### Range expansion correlates with genome size

In all replicates, cells succeed in evolving a cell-cycle to cope with poor nutrient conditions (Fig. 2). Four of the ten replicate experiments (R10, R3, R2, and R8) when ordered by final population size illustrate the following general pattern (Fig. 1; see Fig. S5 for all 10 replicates). On a comparatively short timescale (*t* ~ 10^5^), rapid range expansion to sector 5–7 is observed, while cell density increases in rich environments. Whenever cells adapt to poorer nutrient conditions, higher cell density causing further nutrient depletion can be sustained in previously colonized sectors. Subsequent range expansion occurs in infrequent bursts and is less consistent across replicates than the initial expansion.

**Figure 2:**
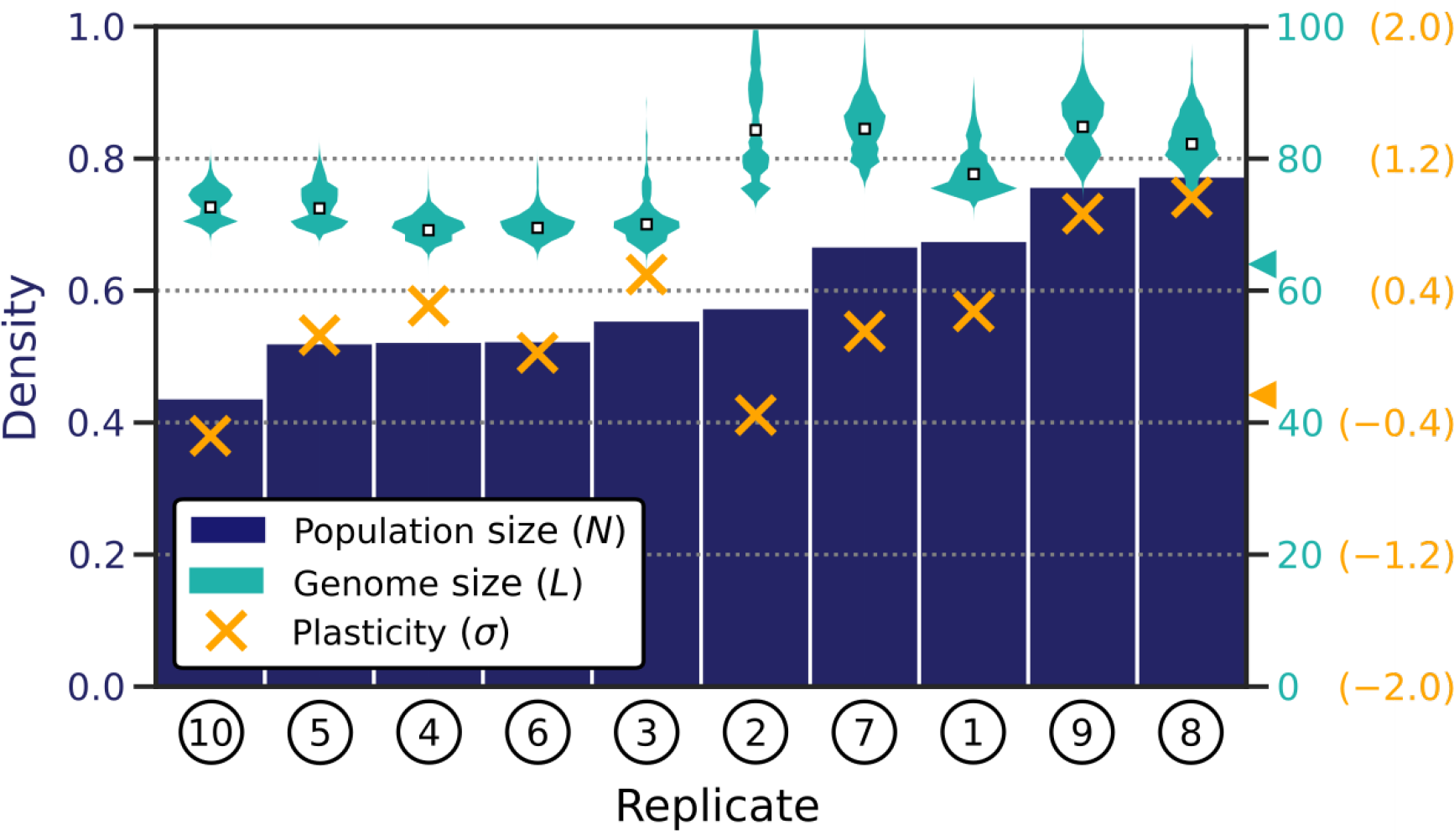
Populations that evolved independently from the same initial conditions show diverse outcomes in population size and average genome size. Replicates are ordered by final population size (dark blue bars). Metrics are averaged across the entire gradient. Plasticity is quantified per cell as 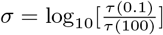. Plasticity of the most common genotype of each sector was measured, and these were then averaged per replicate. Cyan and orange triangles on the right indicate genome size and plasticity for the ancestor.

By the end of the experiment (*t* = 2 . 10^6^), cells have invaded far into the gradient in all replicates and even established themselves at the end of the gradient in some replicates. Persisting at these extremely nutrient-poor conditions requires extraordinary change in cell-cycle regulation. While initial cells obtained S expression once per cycle, the most commonly evolved cell in sector 11 of R8 requires S expression at least 97 times per cycle to complete replication of its genome (*L* = 82, 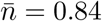).

During range expansion, genome size increases, even though larger genomes require longer to replicate. Coinciding with the early range expansion, average genome size rises from *L* = 64 to *L* ≈ 70. In half of the replicates (e.a. R10 and R3), no further substantial genome expansion occurs. In the other half of the replicates however (e.a. R2 and R8), genomes grow up to *L* ≈ 80 and larger. In R8, there is also transient genome expansion to *L* > 100 along the line of descent. Strikingly, the replicates with the largest genomes reach the greatest population size in the end (Fig. 1, 2). Moreover, genome size is generally larger in nutrient-poor sectors which is remarkable because in those nutrient-poor conditions, selection for shorter genomes would seem to be more severe. Altogether, these observations show that genome expansion is associated with and likely enables adaptation to nutrient-poor conditions.

### Cell-cycle adaptation to nutrient condition

To quantify how individual cells have adapted to distinct and potentially highly fluctuating nutrient conditions along the gradient, we use phenotypic profiling. Single cells are tracked under different fixed nutrient levels, without a gradient or local cell density. During 10^4^ timesteps, we measure in each condition the mean cell-cycle duration *τ*(*n*) and the fraction of cell-cycles that results in proper division *ρ*(*n*), and calculate from these a fitness as 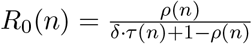.

Evolved cells with larger genomes (R2 and R8) execute faster cell-cycles than cells with smaller genomes (R10 and R3) despite being larger. Provided that the cell does not reach M prematurely (which would decrease *ρ*), faster execution of the cell-cycle is advantageous because it increases the division rate (*R*_0_ increases with decreasing *τ*). Phenotypic profiling thus confirms that genome expansion was adaptive and likely responsible for the previously noted range expansion.

### Individual-versus population-level adaptive strategies

Phenotypic profiling revealed an additional unexpected difference among replicate experiments: different levels of control over the cell-cycle have evolved (Fig. 3). In R3 and R8, the duration of the cell-cycle is modulated by nutrient availability: a longer cell-cycle is executed when conditions are poor, requiring more replication steps. Consequently in replicates where this phenotypic plasticity arises, individual cells demonstrate enlarged viable ranges (e.g. compare cells from R8 with those from R2 in Fig. 3). In R10 and R2, cells did not evolve phenotypic plasticity and even show a slightly negative response to nutrient levels (see Fig. 2).

**Figure 3:**
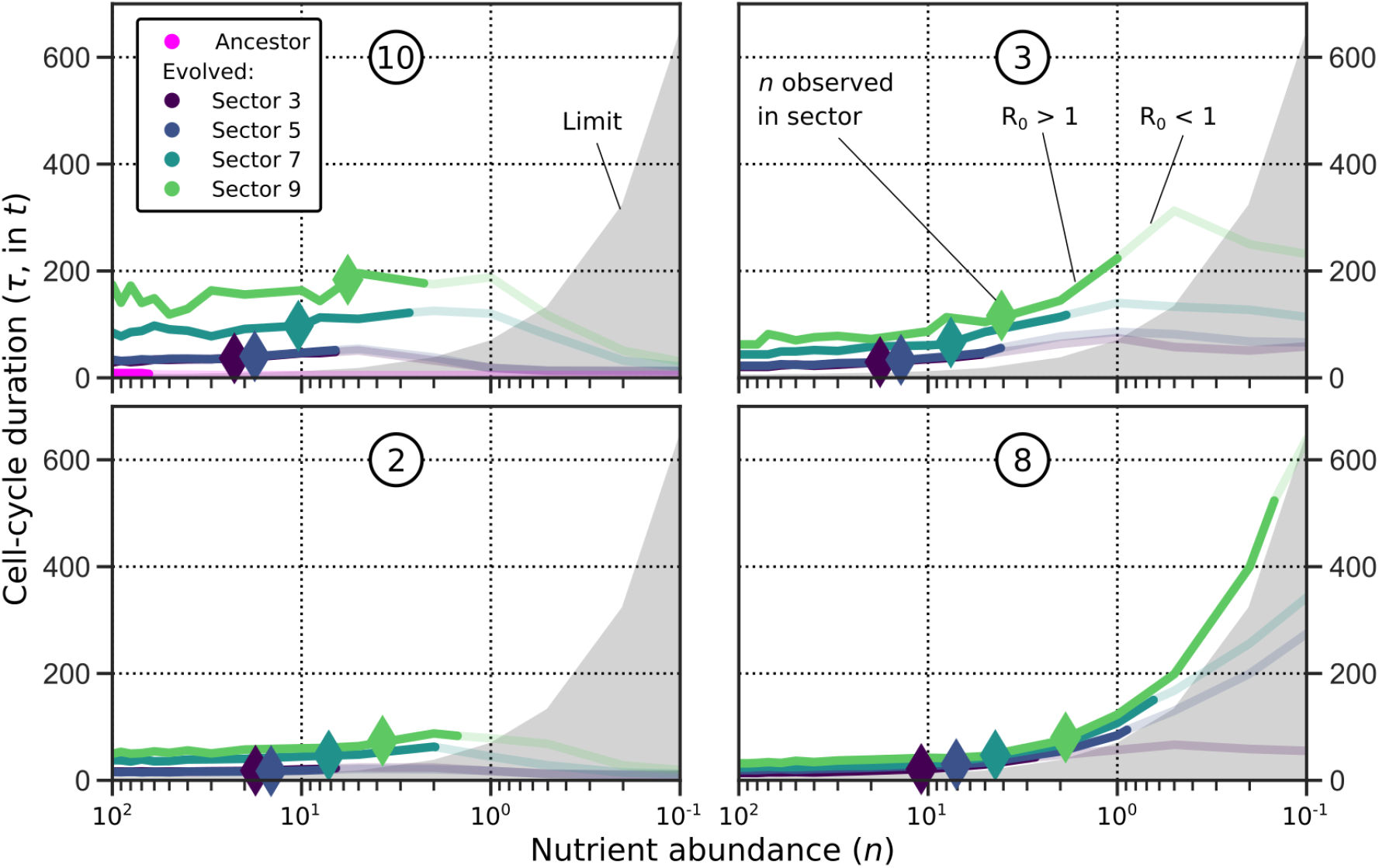
Phenotypic profiling of cells from four populations (R10, R3, R2 and R8). Cells with large genomes (bottom, R2 and R8) regulate faster cell-cycles than those with smaller genomes (top, R10 and R3) in a given sector of the gradient. Because of this, populations in R2 and R8 grow denser, depleting nutrients further (diamonds have moved further to the right). Additionally, cells from R3 and R8 (right panels) time their cell-cycle according to nutrient abundance. This is reflected in the convex nutrient curve and in the larger viable range of individual cells compared to R10 and R2. The grey area marks the minimal duration (*τ*_*min*_) of a complete cell-cycle for a cell with the same genome size as the ancestor (*L* = 64). For all cells, *R*_0_ drops below 1 before *τ* < *τ*_*min*_ because cells have larger genomes (*L* > 64) and because cell-cycle regulation is noisy (*ρ* < 1).

Phenotypic profiling thus reveals two ecological strategies by which populations from different replicate experiments are adapted to the gradient: a generalist strategy in which cells mostly individually tune their cell-cycle to the local conditions, and a specialist strategy in which the population has diversified into cells that are specialised to the different sectors of the gradient. Strikingly, these strategies seem to be independent from the level of genomic complexity in the population (Fig. 2). Evolution has independently produced specialists with small genomes (R10), specialists with large genomes (R2), generalists with small genomes (R3) and generalists with large genomes (R8).

### Evolutionary signatures of generalist and specialist strategies

Further analysis of the replicates showed that the specialist and generalist strategies which were identified in cell-cycle behaviour can also be identified at the phylogenetic and genomic level. The specialist strategy is reflected by a deeper-branching phylogeny than the generalist strategy, as quantified by the *treeness* statistic *T*_*n*_ (Fig. 1; Spearman correlation of treeness versus mean plasticity, *T*_*n*_ × *σ*: *r* = −0.92, *p* = 0.0002, *N* = 10). The treeness shows that specialists form distinct subspecies, whereas generalists resemble a single quasi-species.

At the genomic level, the specialist strategy is reflected by stronger correlation between gene copy number variations and the spatial distribution of cells along the gradient compared to the generalist strategy (Fig. **??**c, *p* = 0.034). In R10 for instance, most cells encode a single copy of gene g6, but cells in sectors 1 and 2 mostly encode two copies.

The same picture emerges when the combined variations in genetic properties such as the activation threshold or regulatory weight are analysed (Fig. S2d, *p* = 0.00044). In sum, generalist and specialist strategies which ultimately lie encoded in evolved genomes and regulatory mechanisms, have an impact on eco-evolutionary dynamics of a population that reaches beyond the cellular phenotype.

### Mechanisms of cell-cycle adaptation to nutrient condition

In our replicate experiments, cells that experienced more genome expansion execute faster cell-cycles, and generalist as well as specialist strategies emerge to deal with varying nutrient conditions. Our modelling framework offers a unique opportunity to investigate how evolved cells accomplish these behaviours mechanistically. To do this, we study cell-cycle regulation at three different levels (Fig. 4). First, dynamic insight into the cell-cycle is provided by the state-space (top panels), which graphs expression states visited during the cell-cycle and the likelihood of transitions between them. These states and transitions are determined empirically by tracking single cells for 10^4^ timesteps (see “Cell-cycle adaptation to nutrient condition”). Second, a concise picture of the regulatory architecture is provided by the network representation (middle panels), which depicts summed interactions between genes. Third, finer regulatory details are provided by the genome representation (bottom panels), which depicts individual interactions between genes and binding sites embedded in the genome. In the next sections, we discuss in terms of these three levels—cell-cycle dynamics, network topology and genome organization—how cells accomplish adaptation to nutrient conditions in different replicates.

**Figure 4:**
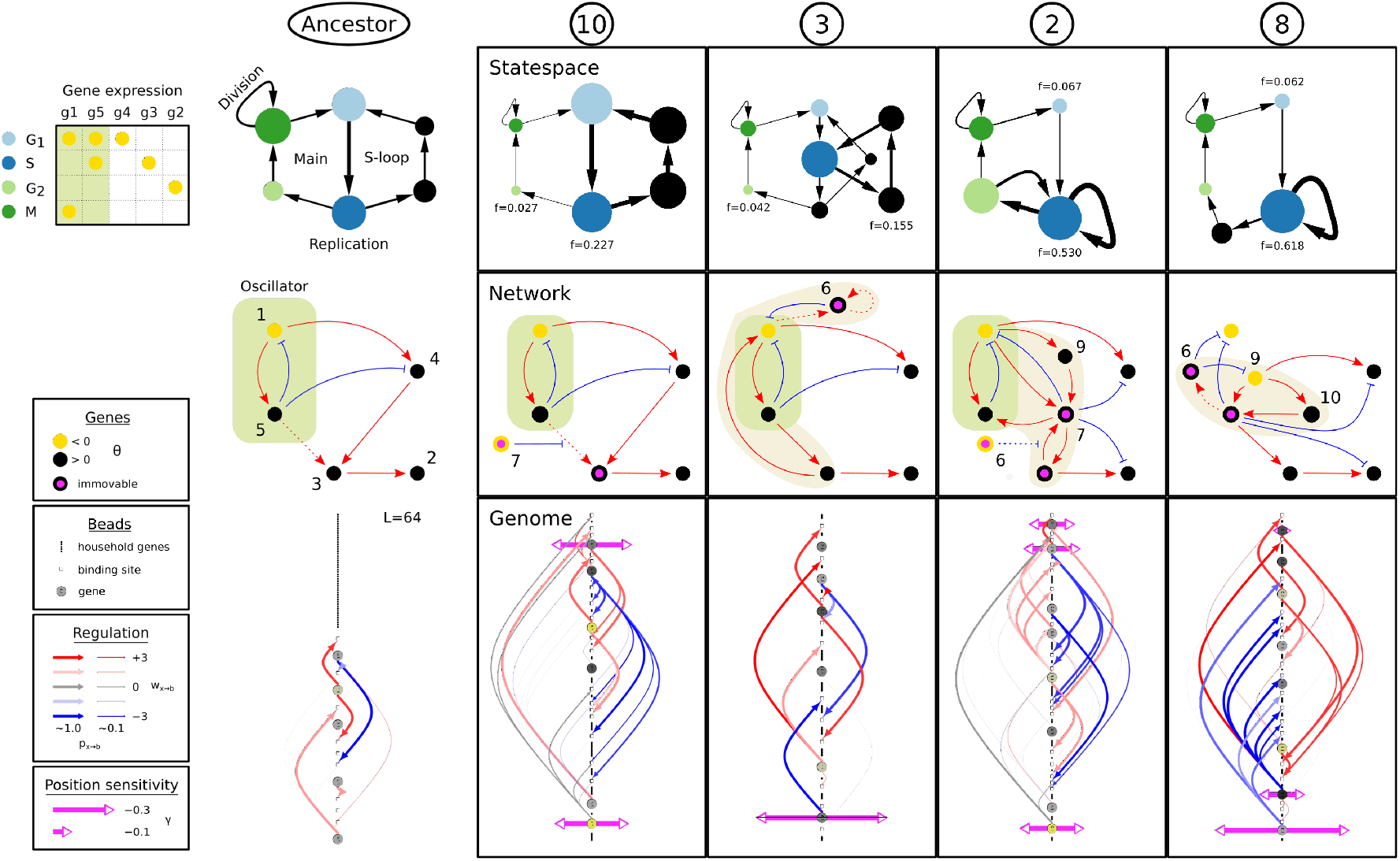
Mechanisms underlying cell-cycle adaptation at three different levels. For a representative from each replicate, the state-space (top panels), network (middle panels) and genome (bottom panels) are shown. The state-space only shows the most frequently observed states and transitions (*f* > 0.02). On the left, the ancestral cell for all replicates is shown, with the central oscillator highlighted in the networks. The additional paler shaded highlight in evolved networks of R3, R2 and R8 identifies other genes in the top level of the regulatory hierarchy. For each replicate, from the 11 most abundant cells per sector a network was chosen that represents the core network (see “Gene family analysis” in Appendix 1).

### Cell-cycle dynamics: contiguous versus intermittent S-phase

For each replicate, state-space reveals how the ancestral cell-cycle was modified by evolution to complete genome replication under limited nutrients, requiring more steps in S before reaching M (Fig. 4, top panels). Besides the original cell-cycle, cells exhibit an alternative cyclic trajectory which runs from S to S and by-passes M. By iterating through this new loop, which we term *S-loop*, cells stall their main cell-cycle and execute additional replication steps.

Differences between the S-loops in state-space reveal how the fitter cells from R2 and R8 manage to execute faster cell-cycles than cells from R10 and R3 (i.e. Fig. 3). In R10, passage through the S-loop takes 4 timesteps, but genome replication only happens in one of these, i.e. in S itself. Thus, cells spend most of their time in non-functional states during stalling of cell division. In R8 in contrast, the S-loop goes directly from S to S, so the cell executes replication contiguously. Under the same conditions, cells from R8 can suffice with an overall much shorter cell-cycle than cells from R10.

### Evolution of the S-phase: rewiring the core oscillator

To understand why a contiguous S-phase evolved in some replicates (e.a. R8) but not in others (e.a. R10), we turn to the underlying regulatory networks (Fig. 4, middle panels). Genome expansion and the resulting complex networks give rise to the more efficient Sloops of R2 and R8. Relative to the ancestor, multiple new genes and interactions arose in R2 and R8, whereas only one new gene arose in R10 and R3. Such genome expansion has resulted in rewiring of the regulatory core of the ancestral network.

In the ancestor, an oscillator composed of g1 and g5 drives both the cell-cycle and the S-loop—the difference between these trajectories is in the activation of g3 by g5 which occurs downstream of the oscillator and hence does not interfere with the cyclic behaviour. The g1/g5 oscillator is still intact in R10, yielding an S-loop with the same period as the main cell-cycle. In R3 and R2, additions to the g1–g5 motif have accomplished shorter S-loops. In R8, the ancestral oscillator has even been entirely replaced by a new module of four genes. Thus, evolution of a contiguous S-phase requires more than genome expansion alone: innovation of regulatory logic is needed to organize an S-loop decoupled from the main cell-cycle. Strikingly, the course of evolution is largely predetermined by early adaptations, as repeating the evolutionary experiments from 2.5 × 10^5^ timesteps onwards generates consistent outcomes (Fig. S6).

## Beyond regulatory logic: cell-cycle timing

The evolved regulatory architecture in each replicate (Fig. 4, middle panels) is conserved throughout the gradient (see “Gene family analysis” in Appendix 1). Specialists and generalists employ this single regulatory blueprint to obtain a wide range of cell-cycle timings across the gradient using genomic speciation and phenotypic plasticity. Through extensive investigation, we could show that network topology and genome organization operate in concert to achieve fine-tuning of the cell-cycle to nutrient conditions (Fig. 4, middle and bottom panels).

### Emergence of functional genome organization

Given the coherence at the network level, variation in cell-cycle timings suggested that other levels such as the genome might be involved. To test this, we performed a gene swap experiment (Fig. 5; Methods). For a given genome, we created mutants which have the location of two genes swapped, for all pairs of genes. Most mutants have reduced fitness compared to the wild-type which can be attributed to 1–4 genes that are located close to the origin or close to the terminus in the wild-type (Fig. 4). In line with this pattern, the genomic location of these genes is more conserved than that of other core genes (Fig. S2b, *p* = 2.6 . 10^−7^).

**Figure 5:**
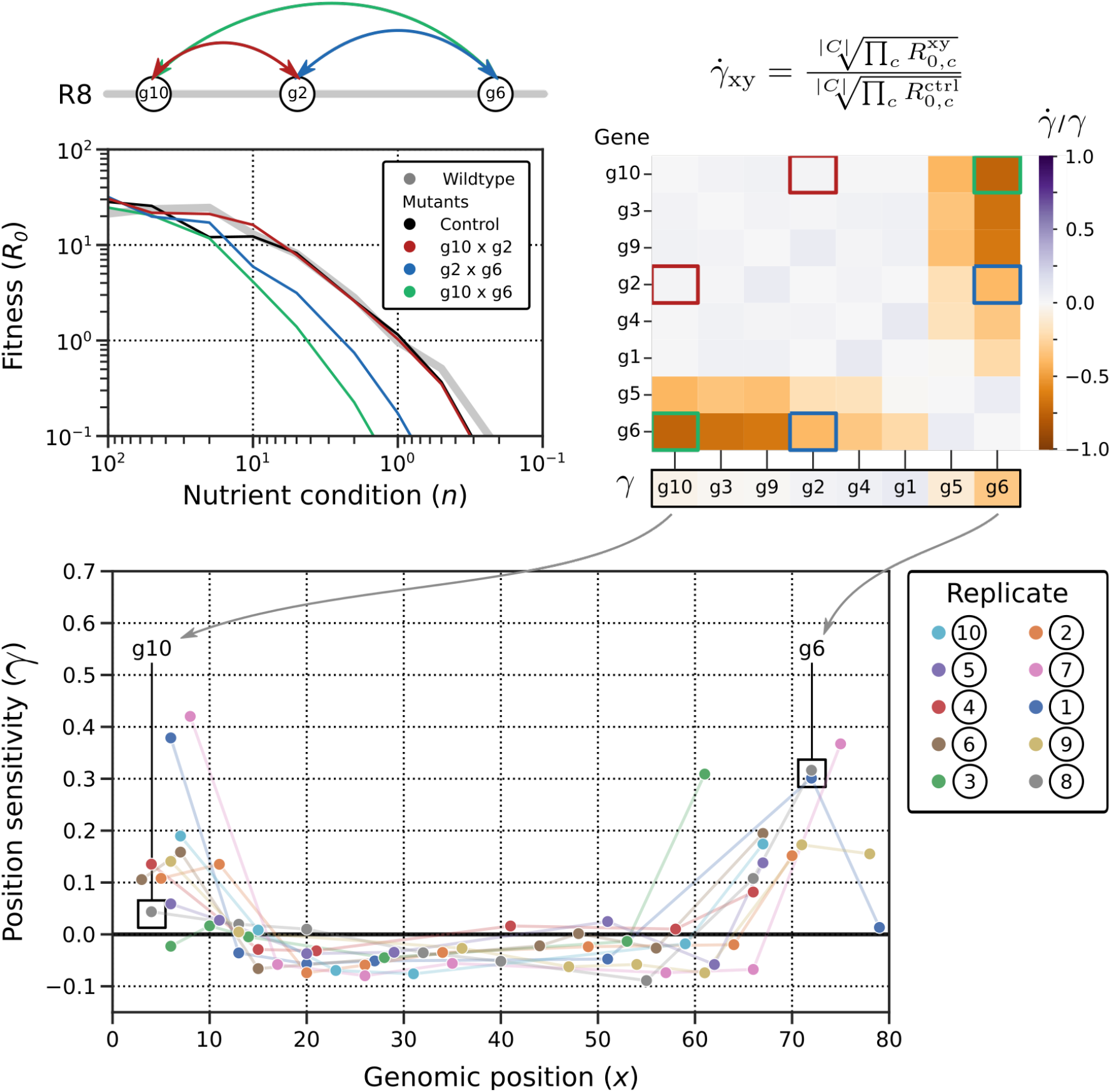
Gene swap experiment reveals that genome organization is functional, in particular for specific genes at the genomic extremities (see text). Top left panel shows the measured *R*_0_-curves for the representative cell from R8 (cf. Fig. 4), the swap control and three swap mutants. The top right shows the matrix with relative fitness of swap mutants compared to the swap control; the three mutants shown on the left are highlighted. From this matrix, relative “position-sensitivity” of each gene is calculated (added row below matrix). In the bottom panel, position-sensitivity is shown for each gene from the 10 representative genomes for R1–10. Each genome harbours at least one gene whose position is crucial for correct cell-cycle behaviour.

Genes that display high sensitivity to genomic location have an important role in the regulatory network: control of cell-cycle progression. They accomplish this control through a weak regulatory interaction that is sensitive to gene dosage. Consequently, such genes link genome organization, which determines when each gene is replicated, to regulatory dynamics, i.e. the likelihood of their interactions. In R8, g6 activation by g5 provides the cue for cell-cycle progression. Since g5 affinity for g6 binding sites is weak, the cell-cycle is stalled in the S-loop long enough, on average, to allow replication to complete. Moreover, the position of g5 and g6 on the genome near the terminus favours g6 activation and progression late in the cell-cycle, when replication is nearly finished. In general, dosage-sensitive genes that need to be active early in the cell-cycle are located near the origin of replication (e.g. g3 in R10), whereas dosage-sensitive genes that need to be active late are located near the terminus (e.g. g5 and g6 in R8). Thus, a simple physical feature like replication-induced change in gene dosage is exploited by an emergent coupling between genome organization and regulatory architecture.

### Generalists: from noisy control to cell-cycle checkpoint

All replicates evolved stochastic control over cell-cycle progression as well as a functional genome organisation to enhance this control. Generalists accomplish phenotypic plasticity through an even tighter coupling between genome and network. The strong response to replication status is achieved by two mechanisms: first, through non-linearity between the cue for progression and progression itself; second, by using gene activation (g6 in R3 and R8) rather than inactivation (g3 in R10, g7 in R2) as the cue. These two features give rise to behaviour that resembles a checkpoint: in the representative cell from R8, cell-cycle progression is nearly four times more likely at the end of replication than it is at the start. Non-linearity between the cue for progression and progression is achieved in at least two ways. In R3 and R8, it presents itself in a timewise manner. The cue for progression, g6 activation, has to be obtained two or three consecutive timesteps in order to steer the cell towards G2. If g6 is only activated once, the cell will revert to the S-loop. As a consequence, when replication increases the likelihood of g6 activation, the likelihood of progression depending on two or three such activations increases quadratically or cubically. Alternatively, in R4, non-linearity arises from the promoter architecture (Fig. 6): activation of g6—which immediately triggers progression—requires binding of g1 and g4 at two different binding sites constituting an AND-gate. When replication increases the probability of each binding event, the likelihood of progression depending on two such events increases quadratically.

**Figure 6:**
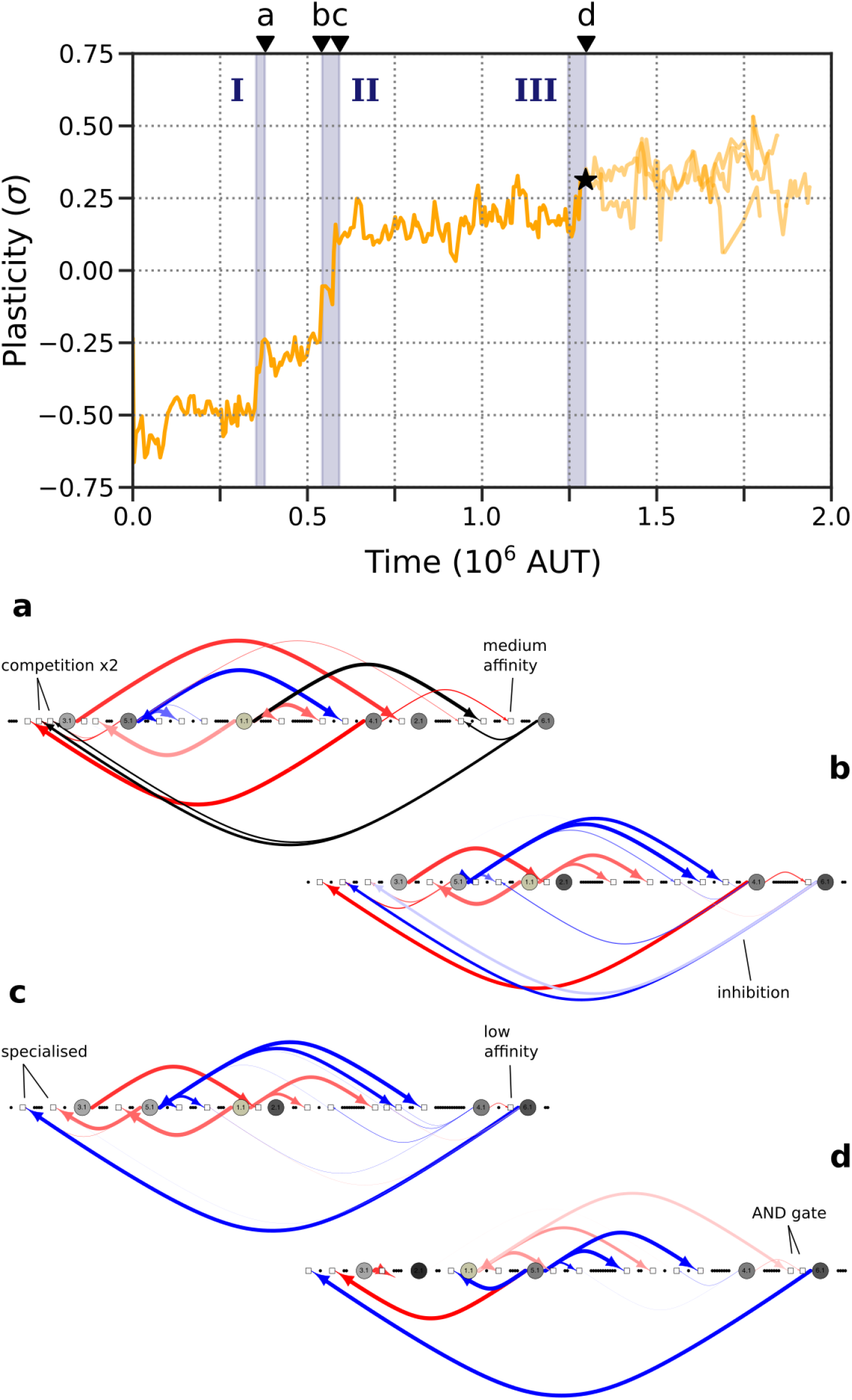
Emergence of plasticity in the generalist replicate R4. Along the ancestral lineage, three phases can be distinguished: I) g6 appears and starts competing with (hindering) g3 activation by g5. II) g6 starts inhibiting g3 directly and the two binding sites of g3 become specialised for g5 and g6. III) g1 starts activating g3 and the AND-gate is formed. Below the time panel, four indicative genomes along the ancestral lineage are shown (birth times indicated by the letters at time points a, b, c, d).

As mentioned, the particular regulatory motif that functions as progression cue (i.e. gene activation or inactivation) also affects how much cell-cycle control can be acquired through genome organization. These observations show that, while genome organization is always relevant whenever low binding affinities play a role, the complex generalist behaviour additionally requires specific network topology to achieve the finer integration between replication and regulation.

### Evolution of a cell-cycle checkpoint

To investigate how the generalist strategy evolved, we focus on one of the replicate experiments, R4, whose checkpoint we mechanistically fully understand as it arose in three separate stages (Fig. 6). The invention of g6, constituting the first stage, improved cell-cycle timing by competitively inhibiting binding of g5 to g3. Plasticity increases because the newly evolved g6 is located at the end of the genome which enhances its regulatory role—promotion of cell-cycle progression. Even more plasticity, i.e. generalism, appears in the second stage, when cells switch to activation as cue for progression. Implicit repression of g3 by g6 becomes explicit through subfunctionalization of two binding sites of g3. In addition, g6 activation by g4 becomes less frequent, replacing binding competition for g3 as timing mechanism. In the third stage, the non-linear response of cell-cycle progression to replication status emerges as g6 activation becomes dependent on weak binding of both g1 and g4. During each stage, reorganization of the genome coincides with (stage I) or rapidly follows (stages II and III) the novel regulatory architecture, underscoring their interplay.

### Specialists: functional genomic speciation

Specialists rely on genomic speciation for adaptation to the gradient. To understand how genomic speciation allows for a wide range of cell-cycle timings, we investigated the previously identified genotypic variation within specialist populations in the context of regulatory topology (see “Gene family analysis” in Appendix 1). Cell-cycle timing is set by the competition between two regulatory genes for a critical target gene (g3 in R10, g7 in R2). Activation of the target gene by an activator (g5 in R10, g3 in R2) leads to stalling of the cell-cycle in the S-loop; inactivation through hindrance of the activator by an implicit inhibitor (g7 in R10, g6 in R2) triggers progression. This competitive balance is tuned to the local nutrient abundance through variation in various genetic properties. For instance, cells in nutrient-poor sectors of R2 encode an additional g7 binding site where competition between the activator and inhibitor takes place. This promotes long cell-cycles, since cell-cycle progression by hindrance of the g7-activator (g3) at all binding sites is less likely. Similarly, short cell-cycles are favoured in nutrient-rich sectors of R10 by means of a second copy of the implicit g3-inhibitor (g7), shifting the competitive balance towards g3 inactivation and cell-cycle progression. Because cell-cycle timing is achieved with a critical noisy interaction, the same regulatory logic can easily be exploited in different conditions by tuning of that interaction. Instead of evolving phenotypic plasticity to cope with different nutrient conditions, specialists increased their evolvability to allow rapid adaptation to those different conditions.

## Discussion

We reformulated a minimal model of the *C. crescentus* cell-cycle to study the evolution of complexity with a focus on gene regulation. The model combines genome evolution and regulatory dynamics with essential cellular dynamics such as death, division, and in particular replication of a quasi-physical genome. Further features were purposely omitted to keep the model maximally insightful. In the model, as in real prokaryotes, replication predominates the cell-cycle yielding an inherent cost to genome expansion. Despite this cost, genomes expanded in our evolution experiments and cells evolved novel regulatory structures to adapt to poor nutrient conditions requiring more complex cell-cycle behaviour. Moreover, to cope with varying conditions, generalist and specialist strategies for cell-cycle timing appeared in different replicate experiments. In both strategies, a single noisy interaction determines cell-cycle timing, establishing efficient utilization of the regulatory repertoire. Generalists especially exploited the positions of the genes involved in this interaction to link replication status (i.e. gene dosage) to regulatory dynamics. The tight integration between network topology and genome organization gives rise to a *de novo* cell-cycle checkpoint. In contrast, specialists employed hard-coded genomic variation of the dosage-sensitive genes to tune cell-cycle timing to different conditions. Thus, range expansion was achieved by plastic individuals or through speciation. The timing strategies and other regulatory mechanisms emerging in replicate experiments demonstrate the model’s success in addressing the evolution of complexity.

### Gene regulation beyond the regulatory network

One of the striking results from our model is the consistent evolution of functional genome organisation in all replicates. Specific genes evolved to be located such that their time of replication aligns with their time of activity, i.e. their role in cell-cycle regulation. Interestingly, similar patterns have been observed in real prokaryotes. In *C. crescentus*, the genes coding for DnaA, GcrA, CtrA and CcrM are positioned sequentially on the genome coinciding with their sequential activation during the cell-cycle (Collier et al., 2007; Seong et al., 2021). In *Escherichia coli*, where the growth cycle dominates replication, transcriptomic data has revealed that the position of many genes correlates with their time of expression during the growth cycle (Sobetzko et al., 2012). Yet even certain binding sites for DnaA in *E. coli* have been shown to play a position-dependent role in cell-cycle regulation (Frimodt-Møller et al., 2017).

Cells couple mechanical progress of the replication fork to cell-cycle control by combining genome organization with specific regulatory architecture. While such feedback from replication on regulation was included in descriptive models of the *C. crescentus* cell-cycle (Li et al., 2008; Shen et al., 2008), our approach has uncovered the mechanisms by which it evolves. The common mechanism was to tune cell-cycle progress by a low-affinity interaction (see “Emergence of functional genome organization”). The probability of such an interaction has not yet saturated which makes it sensitive to replication-induced increase in gene dosage. Another mechanism that generates dosage-sensitivity is competitive inhibition, which evolved in R10, R2, and transiently in R4 (see “Emergence of a cell-cycle checkpoint” and “Specialists: functional genomic speciation”). When genes have similar affinity for a binding site, their relative dosage impacts the outcome of competition, resembling the genetic toggle switch designed by Jaruszewicz-Błońska and Lipniacki (2017). Generalists are prime examples of how evolution succeeded in coupling replication and regulation (see “Generalists: from noisy control to cell-cycle checkpoint”). In R4, cells evolved an AND-gate in the promoter to link gene expression output to two input genes, both of which are amplified by replication (cf. Hermsen et al., 2006). Such motifs may be pervasive in prokaryotes as part of feed-forward loops (Milo et al., 2002). Even more striking is the emergent non-linearity between gene expression and downstream cell-cycle progression that is formed by time integration of the dosage-sensitive interaction, as observed in R3, R8, and R9 (cf. Liu et al., 2015; Malaguti and Ten Wolde, 2021).

It is remarkable that functional genome organization evolved via these mentioned mechanisms even when many complex physical features were not included in the model. Co-operative binding of transcription factors to DNA (Hermsen et al., 2006) or regulation beyond the level of transcription (Bryant et al., 2014; Frimodt-Møller et al., 2017; Krogh et al., 2018) would only seem to provide more routes for evolution to couple replication to regulation. In *C. crescentus*, chromatin states are used to control gene expression during the cell-cycle (Collier et al., 2007; Seong et al., 2021). Shortly after the transcription factor DnaA initiates replication, the *dnaA* locus near the origin is replicated yielding two hemi-methylated promoters. As DnaA is preferentially transcribed from a fully methylated promoter, the change in chromatin state leads to collapse of expression preventing reinitiation of replication. For CtrA, replication has the opposite effect. CtrA is preferentially transcribed from a hemi-methylated promoter, so expression starts when the *ctrA* locus is replicated. These examples illustrate that prokaryotes have complex biochemistry available, which can be used to form an even better coupling between replication and regulation than sheer dosage changes.

We have demonstrated that integration between genome organization and network topology can evolve in many ways. Yet biochemistry and biophysics inherently couple replication to regulation—evolution merely exploits this link. Fluctuations in gene copy numbers naturally lead to fluctuations in transcript numbers (Walker et al., 2016). While fluctuations are largely buffered on the protein level, this most likely represents an evolved state as many cellular processes do not benefit from being driven by the cell-cycle. For instance, *Synechococcus elongatus* has evolved additional regulatory mechanisms to decouple its diurnal cycle from its cell-cycle (Paijmans et al., 2016). Thus, interaction between replication and regulation should be the default expectation in prokaryotes.

### Evolution of complexity

In our replicate evolution experiments, cells adapted to new conditions through an increase in cellular complexity. Besides expansion of the regulatory repertoire, genome organization evolved to improve cell-cycle timing to different external conditions. By including both genome structure and network topology, the interaction between these layers itself evolves, representing a new dimension of complexity that gives rise to “no-cost generalism” (Remold, 2012): generalists do not encode more regulatory components, but organize their components in a specific way. Because reorganization of the regulatory network is difficult, the emergence of this type of complexity is contingent on the right motifs appearing early in evolution which explains why it is not a ubiquitous outcome. Yet, between replicates, generalists appear to have adapted most successfully, exemplified by the two replicates with the largest final populations, R8 and R9. Thus, the outcomes of our model lend support to the theory that generalism is a stable strategy and not just an evolutionary intermediate state (Dennis et al., 2011; Loxdale et al., 2011; Sriswasdi et al., 2017).

Most substantial adaptation was due to the expansion of genomes and their encoded regulatory networks, forming novel regulation to organize replication more efficiently in an S-phase. Previous models also observed network expansion (Soyer and Bonhoeffer, 2006; Cuypers and Hogeweg, 2012, 2014), but with mostly non-adaptive forces implicated. In these models, the cost to complexity was relatively low (Soyer and Bonhoeffer, 2006) or absent (Cuypers and Hogeweg, 2012, 2014), allowing network expansion to accommodate weak selection for robustness or evolvability. In our case, the energetic constraint on genome size limits such complexity (see Fig. S4). To our knowledge, this is the first model that demonstrates the role of adaptive forces for the evolution of complex regulation. Although adaptive forces have perhaps been historically implicated, they have never been computationally tested. What stands out from this and previous theoretical work is that an increase in complexity is possible under very different evolutionary regimes, each of which leaves a potentially characteristic signature on the evolutionary dynamics and on genome and network organization. This brings us one step closer to a theoretical framework for the evolution of complexity that will help us understand how life ventured beyond the first replicators, to prokaryotes, eukaryotes, multicellularity, and sociality.

## Methods

We study the evolution of regulatory networks in an individual-based model embedded on a grid. Each cell in a population consists of a genome with regulatory genes (or genes, for short) and binding sites, that together form a regulatory network. Expression of five core gene types (g1–5) defines the cellular state 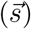. Reproduction takes place when a cell completes a trajectory through four states that represent the *Caulobacter crescentus* cell-cycle (taken from the model of Quiñones-Valles et al., 2014; Sánchez-Osorio et al., 2017).

### The cell-cycle

Quiñones-Valles et al. (2014) derived from literature a Boolean network for cell-cycle regulation in the model bacterium *C. crescentus*. With five core regulatory genes (CtrA, GcrA, DnaA, CcrM and SciP), the network already yields a cyclic attractor of four states, resembling a cell-cycle. We use these states—which we will denote G1, S, G2 and M—to define the cell-cycle in our model.

When a cell acquires M expression, one of two things happens. If the cell has progressed through all previous stages, it divides. A new cell is formed at an adjacent site on the grid, killing if necessary the cell that occupied that location. If the cell reaches M prematurely however, it dies. For proper cell-cycle progression the order of stages is important, but they do not have to follow on one another directly. A cell in G1 can reach any number of alternative states or even perform the expression that defines G2. This merely stalls its cell-cycle until it acquires the expression corresponding to the next stage (S in this case).

One of the key and most time-consuming events in the prokaryotic cell-cycle is replication of DNA. The explicit incorporation of genome replication in our model leads to a natural constraint on genome size: an information cost. Cells replicate a piece of their genome (*n* beads, where *n* is the nutrient abundance) every timestep they spend in stage S. The copied beads participate in the regulatory dynamics. With time spent in S, the copy number of each gene increases from one to two, impacting the regulatory dynamics (see next section).

Initially, we evolve individuals with the basic cell-cycle circuit under unlimited nutrients (*n* = ∞), such that the entire genome is replicated in a single step (see “Evolution of *C. crescentus* cell-cycle under stochastic gene expression”). This allows cells with the Quiñones-Valles et al. (2014) regulatory network, which spend only one step in S, to successfully complete their cell-cycle. Afterwards, we expose cells to nutrient limitation, which requires them to express multiple replication steps (S) in one cell-cycle (see Results). Cells are inoculated on a nutrient gradient (*n_tot_* = [2, 80]) and nutrients are locally depleted by other cells, so that a site with *x* neighbours has a nutrient level 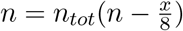.

Due to real-time genome replication and nutrient limitation, there is a strong cost to genome size. Cells with a smaller genome require fewer replication steps, which allows them to complete their cell-cycle faster and produce more offspring. Note that regulating more replication steps than required (i.e. when the genome is already completely replicated) does not harm the cell except that it costs time, decreasing reproduction rate. Besides regulatory components, cells need to maintain 50 household genes in their genome. When a cell-cycle is completed, the reproduction probability is reduced by 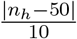 where *n_h_* is the number of household genes.

### The genome

Genes, binding sites and house-hold genes are organised on a linear beads-on-a-string genome with an origin of replication and a terminus at the two opposite ends (cf. Crombach and Hogeweg, 2007). A gene is expressed if the sum of regulatory effects at its upstream binding sites reaches its specific activation threshold *θ*:

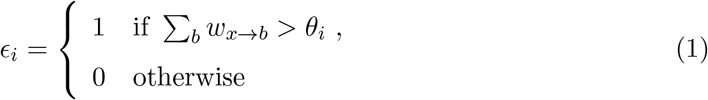

The regulatory effect *w*_*x*→*b*_ results from the binding of expressed gene *x* to binding site *b*. This effect depends on the regulatory weight of the binding site and the bound gene:

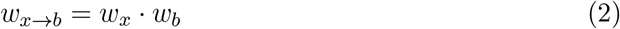

Only one gene may bind per binding site, and this is (re-)determined stochastically every timestep. All genes that are expressed at that moment have a probability to bind, based on their affinity for the binding site. There is always a probability that no gene binds, the relative propensity of which is set to 1:

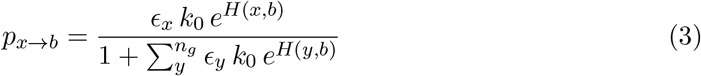

Here, *H*(*x, b*) is the Hamming distance between the binding sequences of *x* and *b* (bitstrings of length *l*_*B*_ = 20) and *n*_*g*_ is the number of genes in the genome. The parameter *k*_0_ was set to *k*_0_ = 1.0 . 10^−7^ because this leads to low affinity between two random sequences on average, strong binding for two very similar sequences (*p*_*x→b*_ → 1.0), and an intermediate region where affinity can increase or decrease in a graded fashion between these two extremes.

### Mutations

Mutations happen at the level of the genome. Upon division of a cell, genes, binding sites, and household-genes in the new daughter cell can undergo various mutations.

First, the genetic properties of beads can be mutated. With probability *μ*_*B*_ per position, a bit in the binding sequence 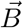 is flipped. With probabilities *μ*_*w*_ and *μ*_*θ*_ per bead, the regulatory weight *w* and the activation threshold *θ* are mutated, respectively (*μ*_*θ*_ only applies to genes). The value of *θ* or *w* is then incremented or decremented (probability 0.8) or randomly redrawn from the range [−3, 3] (probability 0.2).

Second, each bead has a probability to be duplicated (*μ*_*dup*_), deleted (*μ*_*del*_) or relocated (*μ*_*rel*_). The new location of a duplicated or relocated bead is random. When genes are duplicated, deleted or relocated, adjacent upstream (“proximal”) binding sites move with them.

Third, there is a per-genome innovation rate for binding sites (*μ*_*b,in*_) and for genes (*μ*_*g,in*_). These result in the creation of a new bead of the respective type with random properties and at a random location in the genome.

As mentioned, the cellular state 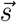 is defined by the expression of the five genes originally modelled by Quiñones-Valles et al. (2014). We do not want to constrain the regulation of these transcription factors themselves during *in silico* evolution. Therefore, we assign gene types *t* to unique combinations of binding sequence 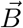 and regulatory weight *w* (these parameters define the gene product; in contrast, *θ* is a property of the locus) where g1–5 denote CtrA, GcrA, DnaA, CcrM and SciP. Genes keep their identity even if 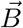 or *w* changes. New gene types are only assigned upon gene innovation or when a duplicated gene diverges from the other copy and establishes a new unique combination of 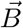 and *w*.

### Gene swap experiment

A gene swap experiment was used to assess the importance of genome organisation and to pinpoint genes whose position is crucial to cellular fitness. First, a wildtype genome is constructed from the original by placing all binding sites directly adjacent to their down-stream gene. Then, genes are spaced from one another such that the maximal number of binding sites of any gene can fit in between. In the resulting wildtype, genes together with their binding sites can be swapped without displacing/moving any of the other genes in the genome. For each pair of genes, a mutant is created with the genomic location of those two genes swapped. Then, in addition to the wildtype, the fitness of each of these mutants is estimated in a range of fixed nutrient conditions *C* = {100, 50, 20, 10, 5, 2, 1, 0.5, 0.2, 0.1}:

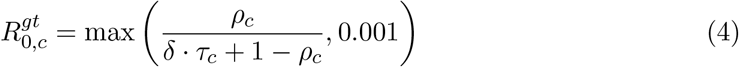

Here the genotype *gt* stands for the wildtype or any mutant *xy* with genes *x* and *y* swapped. The fitness of each mutant across all conditions relative to the wildtype is calculated by comparing the harmonic mean:

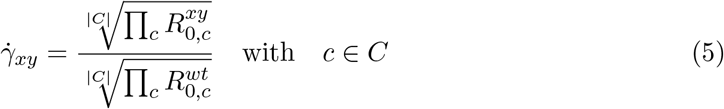

Then, to find the relative sensitivity to relocation of a particular gene, its contribution to the fitness reductions in all its swaps is calculated. Basically, the off-diagonal *xy*-elements in the matrix *γ* are normalised by the average of the column (all contributions of gene *y*), and then averaged per row (all contributions of gene *x*):

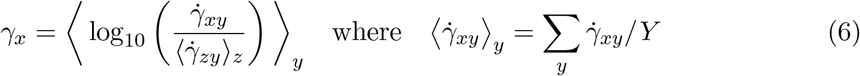

**Table 1:**
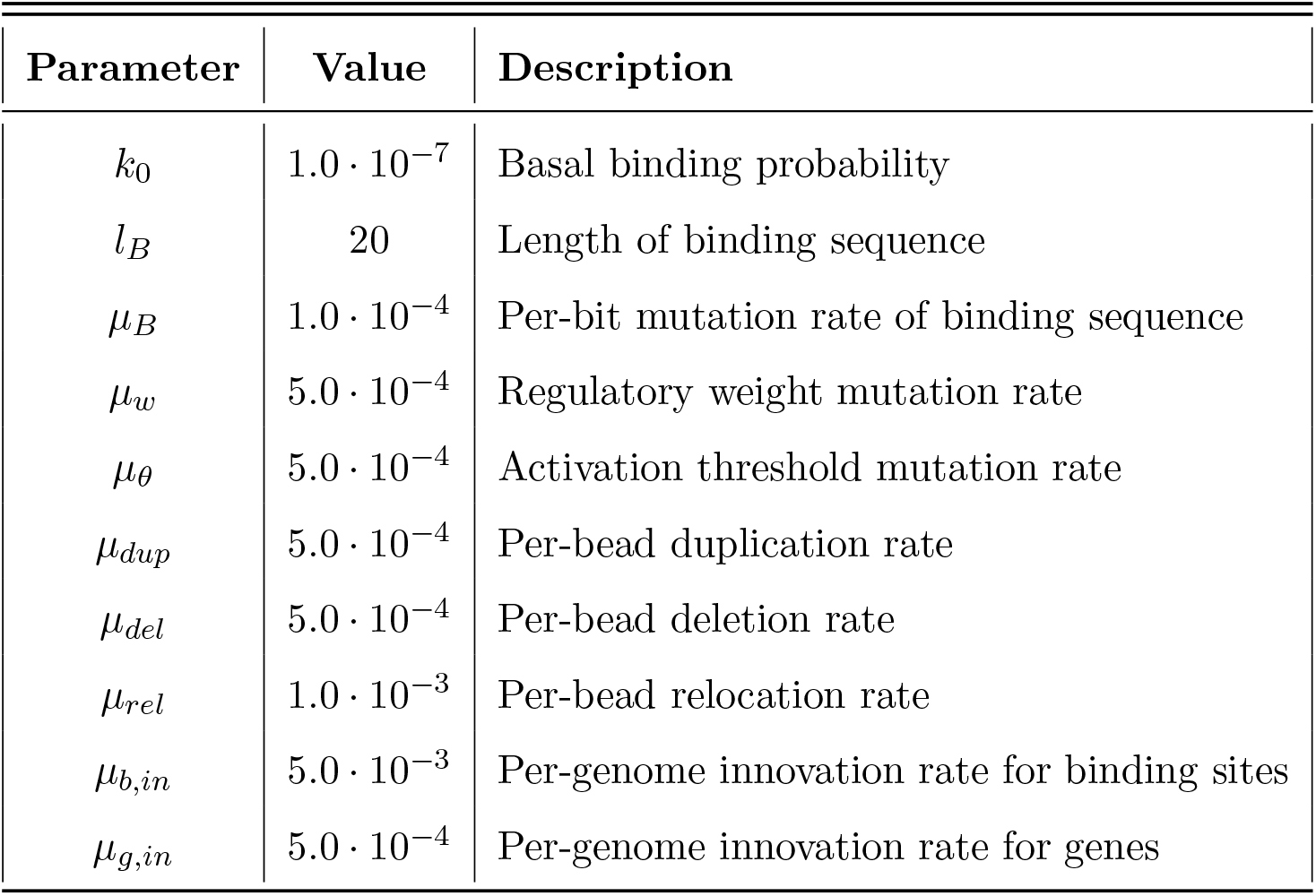
Parameter values used in the evolution experiments

| Parameter | Value | Description |
| --- | --- | --- |
| $k_0$ | $1.0 \cdot 10^{-7}$ | Basal binding probability |
| $l_B$ | 20 | Length of binding sequence |
| $\mu_B$ | $1.0 \cdot 10^{-4}$ | Per-bit mutation rate of binding sequence |
| $\mu_w$ | $5.0 \cdot 10^{-4}$ | Regulatory weight mutation rate |
| $\mu_\theta$ | $5.0 \cdot 10^{-4}$ | Activation threshold mutation rate |
| $\mu_{dup}$ | $5.0 \cdot 10^{-4}$ | Per-bead duplication rate |
| $\mu_{del}$ | $5.0 \cdot 10^{-4}$ | Per-bead deletion rate |
| $\mu_{rel}$ | $1.0 \cdot 10^{-3}$ | Per-bead relocation rate |
| $\mu_{b,in}$ | $5.0 \cdot 10^{-3}$ | Per-genome innovation rate for binding sites |
| $\mu_{g,in}$ | $5.0 \cdot 10^{-4}$ | Per-genome innovation rate for genes |

## Acknowledgements

The authors gratefully acknowledge the help of Jan Kees van Amerongen for running the local computer cluster, Bram van Dijk for his advice during writing of the model’s code, and Rutger Hermsen for his advice on a suitable physical function for the binding probability between a gene and a binding site.

