## Supplemental Material for "Evolution of Regulatory Complexity for Cell-Cycle Control"

July 28, 2021

### Appendix 1: Supplementary Results

#### Evolution of *C. crescentus* cell-cycle under stochastic gene expression

First, the network topology of Quiñones-Valles et al. (2014) was implemented in our modelling framework. Each of the five transcription factors was given a sufficiently unique bitstring—reassigning the sequences to genes in a different way did not change the results. For each interaction between two genes, a separate binding site was placed upstream of the target and with the bitstring matching to the regulating gene. With this genotype, an evolutionary experiment was performed without explicit replication, i.e. without the need to modify cell-cycle behaviour. Note however that the original state-space as shown in Quiñones-Valles et al. (2014) contains two attractors: one is the cell-cycle and another is a seemingly non-functional point attractor.

The most common genotype after $10^5$ timesteps was used for the evolutionary experiments described in the main text. Although average gene copy number and binding sites increased during the experiment, the most common genotype was streamlined, only retaining a subset of the original interactions. At the functional level, such streamlining appeared to be robust: the most common genotype of four more replicate experiments revealed the same network topology. In some cases, and in fact in most genomes of each population, more genes appeared, although without a substantial functional role.

One crucial improvement is the appearance of baseline activity of g1 ($\theta < 0$). This abolishes the second non-functional attractor that was present in the network of Quiñones-Valles et al. (2014) and which has become a liability in our model due to the stochasticity in regulatory dynamics. Another notable change is the appearance of the activation of g3 by g5. Although this change likely did not alter the cell-cycle in this simple initial experiment, it became an important subject for evolution in the ten replicate evolution experiments (see main text, Fig. 4).

### Gene family analysis

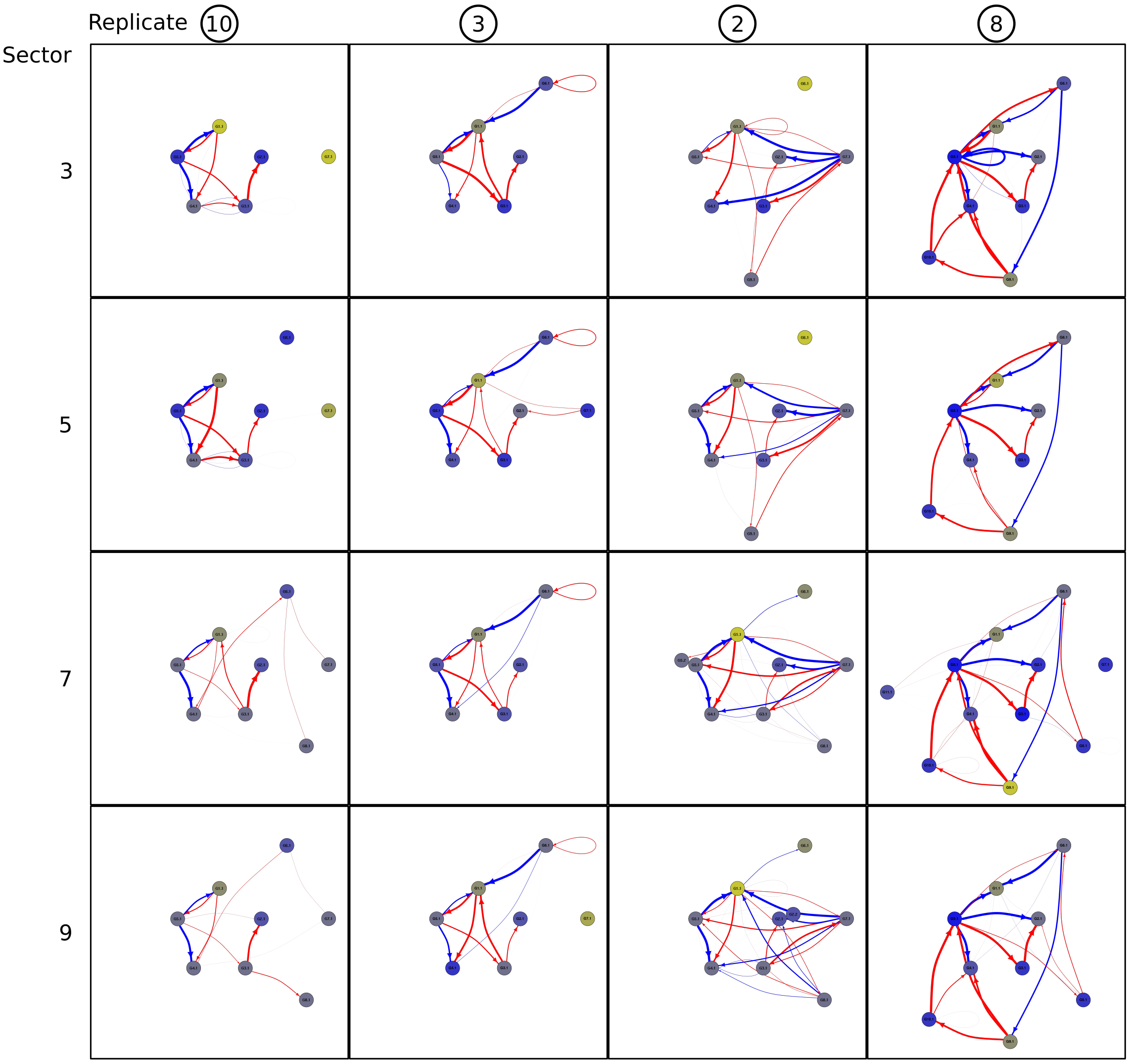


Figure S1: The architecture of gene regulatory networks varies between replicate evolution experiments but is comparable within each experiment. While some variation across the gradient is functionally relevant (e.g. the extra gene g8 in sector 9 of R8), most variation is likely neutral (e.g. the two extra genes g7 and g11 in sector 7 of R8, neither of which was observed to be expressed during phenotypic profiling for over 1500 timesteps).

To assess genotypic variation in populations, and study specific patterns in that variation (i.e. Fig. S1), we performed a gene family analysis (Fig. S2, S3). We defined gene orthologies per population and assigned to these the genes from all genomes in that

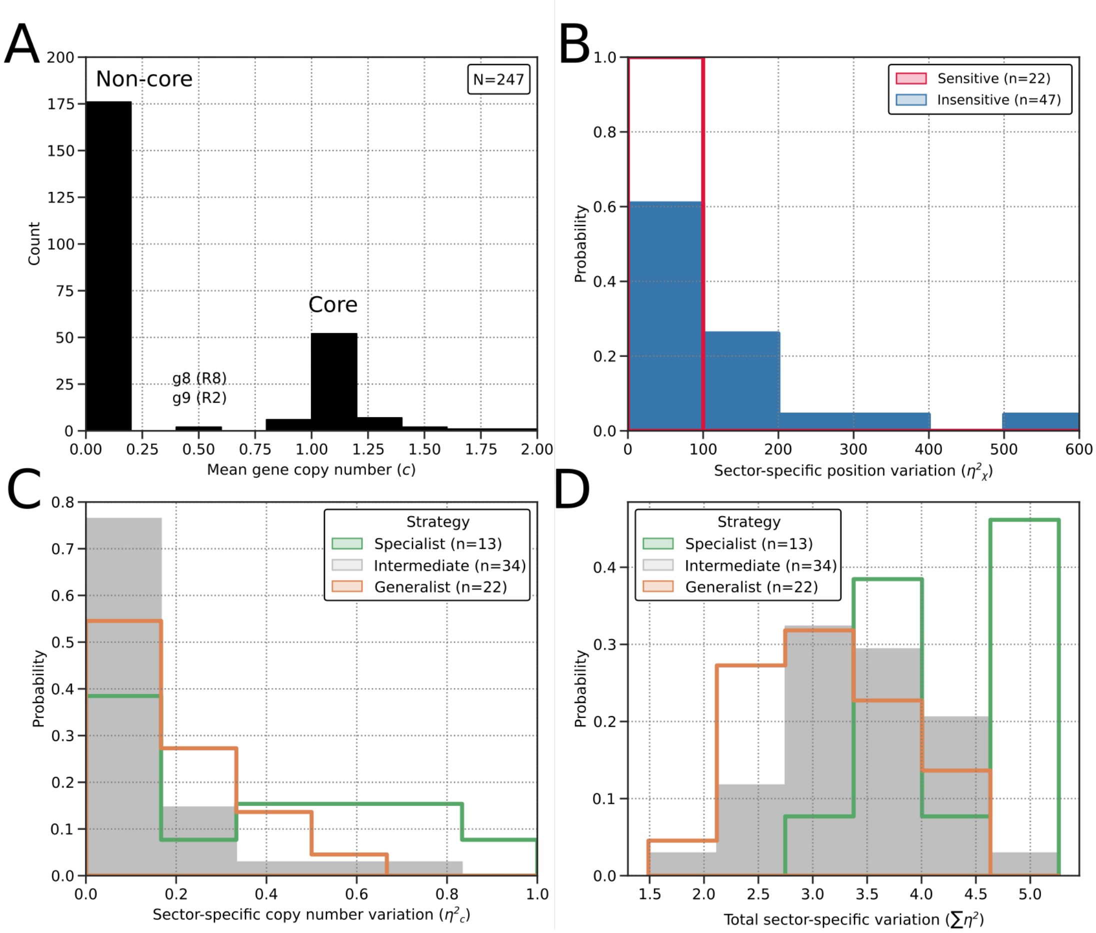


Figure S2: Signatures of function and strategy in core gene families. (a) Each population has an easily identifiable core set of gene families for which mean copy number is centered around 1.0. (b) Genomic location varies less across sectors in dosage-sensitive genes than in other core genes (Mann-Whitney $U = 134$, $p = 2.56 \cdot 10^{-7}$; see "Emergence of functional genome organization"). (c & d) Core gene families of specialist populations are more specialised to the gradient than families of generalist populations. For each family is shown (c) sector-specific variation in copy numbers (generalist × specialist: Mann Whitney $U = 89$, $p = 0.0339$), and (d) sector-specific variation in 9 other genetic parameters combined (see Methods; generalist × specialist: Mann Whitney $U = 45$, $p = 4.36 \cdot 10^{-4}$). Based on the level of phenotypic plasticity (see main text, Fig. 2), R3, R8 and R9 are annotated as generalist ($\sigma > 0.4$); R2 and R10 are specialist ($\sigma < 0.0$); the five remaining replicates are intermediate ($0.0 \leq \sigma \leq 0.4$).

population. For each population, we find a small set of core gene families (mean copy number per cell $> 0.95$ and present in $> 95\%$ of unique genomes), which always include g1–5 (Fig. S2a). Out of the non-core families, the large majority is only found in a few genomes (mean copy number per cell $< 0.1$, present in $< 10\%$ of unique genomes), which accounts for less than 2% of gene content on average. Two interesting exceptions are g8 in R8 and g9 in R2, which are ubiquitous in one half of the gradient and completely absent in the other (mean copy number $\approx 0.5$, see Fig. S2a). Core gene families do not show such sharp presence-absence variation across the gradient, but some of them vary in

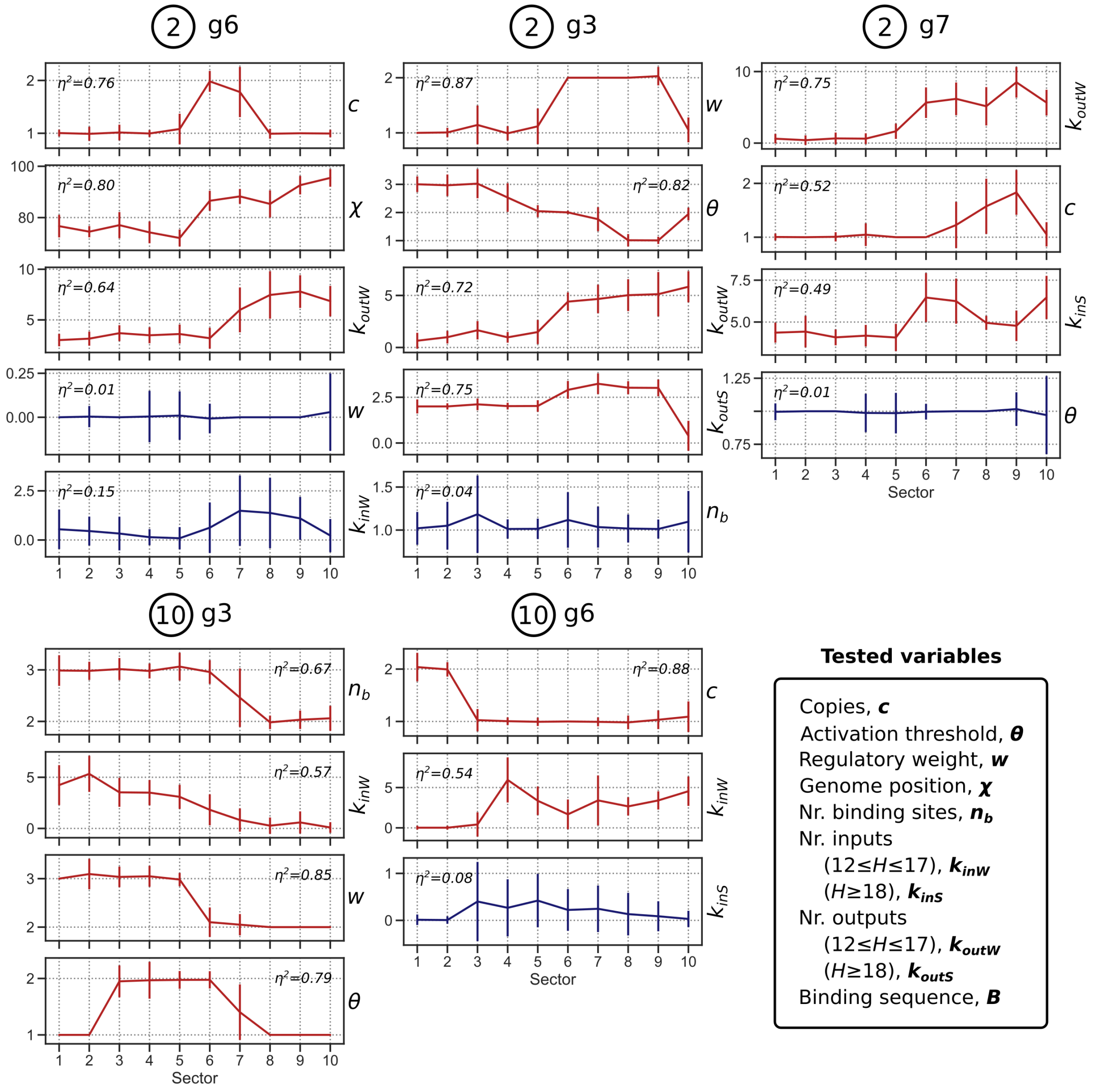


Figure S3: Core timing genes of specialists (R2 and R10) are specialized to sectors of the gradient in multiple ways. For each dosage-sensitive gene pinpointed by the gene swap analysis, sector-specific variation is shown for properties where the gene ranks high (top 10; red lines) or low (bottom 10; blue lines) in terms of $\eta^2$ among all 69 core gene families. The number of occurrences of high ranks for dosage-sensitive genes of specialists is far greater than the expected value ($X = 16$, $E(X) = 6.73$), whereas the opposite is true for dosage-sensitive genes of generalists ($X = 3$, $E(X) = 9.62$).

copy numbers. By comparing variation between sectors to variation within sectors (using ANOVA's $\eta^2 = \frac{SS_{groups}}{SS_{groups}+SS_{errors}}$), we can quantify several types of specialization to the gradient. Besides copy number specialization, we obtain $\eta^2$ for 9 other properties of the 69 core gene families: activation threshold $\theta$, regulatory weight $w$, position on the genome $\chi$, number of binding sites $n_b$, regulatory input ($k_{inS}$ and $k_{inW}$, for edges with strong and weak affinity, respectively), regulatory output ($k_{outS}$ and $k_{outW}$) and binding sequence $B$ (see Fig. S3).

Figure S2c,d shows the degree of specialization ($\eta^2(c)$ and $\sum \eta^2$) in gene families from all populations subdivided by strategy. Gene families from specialists tend to be more specialized to the gradient than families from generalists (Fig. S2). In Figure S3, we show examples of how specialists adapt to the gradient through genomic differentiation. This analysis shows that the genotypic structure of a population recapitulates its eco-evolutionary strategy (see main text).

Several more patterns can be found in the genotypic data. As expected, the number of binding sites upstream of a gene correlates with the number of regulatory inputs ($n_b \times k_{inS}$: $r = 0.84$, $R^2 = 0.71$, $p < 0.0005$, $N = 247$). Similarly, the sector-specific variation of these properties is also correlated ($\eta_n^2 \times \eta_k^2$: $r = 0.7$, $R^2 = 0.49$, $p < 0.0005$, $N = 247$). Interestingly, specialization of a gene's binding sequence is negatively correlated with the number of binding sites that it targets ($n_b \times k_{inS}$: $r = -0.66$, $R^2 = 0.44$, $p < 0.0005$, $N = 247$). Thus, in line with literature, the binding domain of a transcription factor is more constrained by purifying selection if it coevolves with more targets.

### Model testing

To assess how different features of our model impact its dynamics, we performed evolution experiments with modifications of the main model (Fig. S4). First, the constraint on genome size is removed by letting the nutrient level $n$ define the proportion of the genome that is replicated in one timestep instead of the absolute number of beads. Under such proportional replication, massive and rapid genome expansion occurs. Although most added beads likely appear due to loss of purifying selection, population size increases more rapidly than in the standard model, showing that genome expansion brings about adaptation. The 10 replicates with proportional replication were terminated early because too large genomes become a computational burden.

In constrast, when stochasticity in binding dynamics is removed—i.e. at each binding site, interactions with genes with an affinity $H \geq 17$ are averaged over time—genome expansion, and thereby adaptation, are entirely absent. Instead, genomes are streamlined very early from $L = 64$ to $L \leq 60$ so that the short ancestral cell-cycle—which, in the absence of stochasticity, is at the theoretical optimum for high nutrient levels—can be employed in a slightly wider niche range. In 3 out of 10 replicates, cells expanded their range even further by loss of 8 household genes (obtaining $L = 52$) despite large fitness penalties.

The above two modifications of the model demonstrate how crucial selection is for

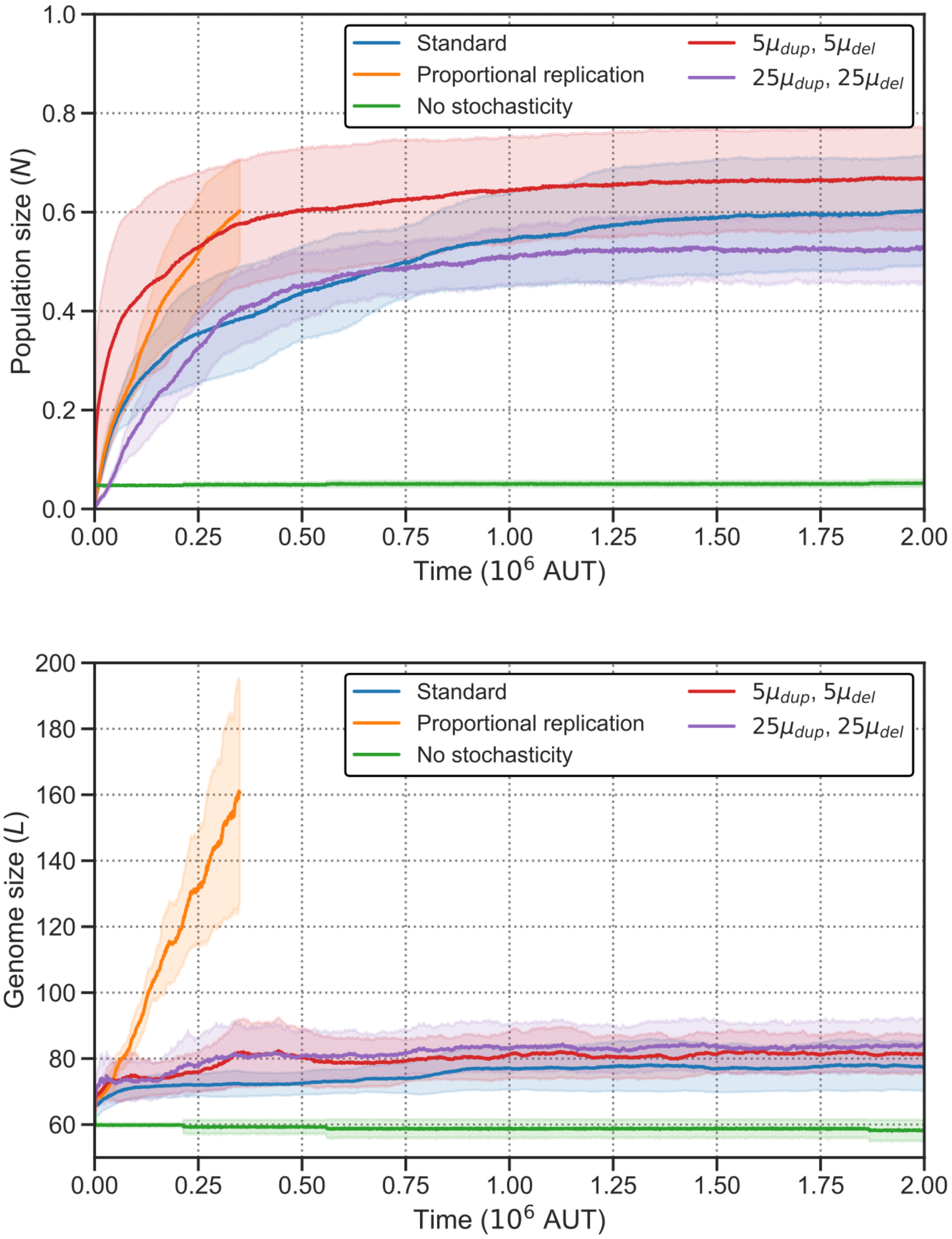


Figure S4: Testing of model features. (a) Population size and (b) genome size versus time, averaged over multiple replicate evolution experiments for different model modifications ($n = 5$ for the two models with higher duplication and deletion rates; $n = 10$ for the other three models). When replication time does not depend on genome size, massive genome expansion ensues. Conversely, without stochasticity in binding dynamics, no adaptation or genome expansion occurs. Increased duplication and deletion rates have a milder impact on evolutionary dynamics.

genome size evolution. Without altering the mutation rates (specifically duplication, deletion and innovation), we obtain two extremes in genome size evolution. When we increase the rate of duplication and deletion 5-fold, genome expansion also becomes more prevalent, but the energetic cost of replication still dominates and generally prevents the emergence of genomes larger than $L \approx 80$. Yet, the greater influx of additional regulatory elements helps to reorganize cell-cycle regulation and thus favours adaptation, as seen in the greater population sizes than the standard model during the entire experiment. There actually appears to exist an optimum for the rate of duplication and deletion, as another 5-fold increase (25-fold compared to the standard model) yields smaller populations than the standard model.

### Appendix 2: Supplementary Figures

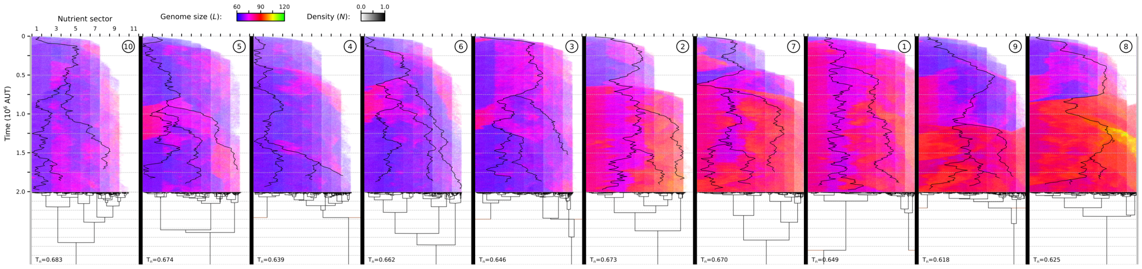


Figure S5: Evolutionary dynamics of all ten replicate experiments, ordered by final population size (see main text, Fig. 1).

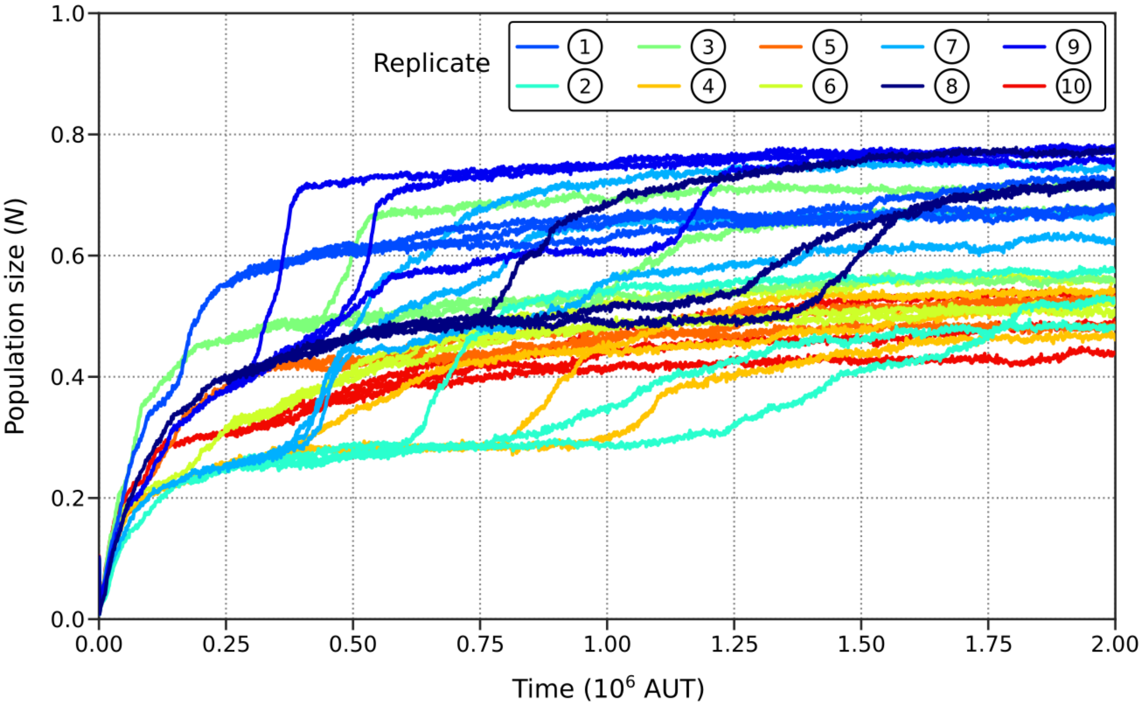


Figure S6: Adaptation becomes constrained very early in evolution, as shown by population growth of the ten replicate evolutionary experiments plus two alternative trajectories starting at $t = 2.5 \times 10^5$ time steps (lines with same color as original experiment). Alternative trajectories generally end up with similar population size (blue lines high, red lines low), showing that adaptive potential is already constrained by $t = 2.5 \times 10^5$.
